# Genome-Level Hierarchical Attention Transformer with Multi-Head Attention Weighted Sum for Broad-Spectrum Antimicrobial Resistance Prediction and Discovery of Resistance-Related Genomic Contexts

**DOI:** 10.64898/2025.12.22.695895

**Authors:** Sein Park, Hyounggyu Kim, Kihyun Lee, Hyun-Seok Oh

## Abstract

Antimicrobial resistance is a growing global health concern, requiring reliable tools for predicting resistance across a wide range of bacteria and antibiotics. In this study, we introduce a genome-level hierarchical attention transformer (GL-HAT) that integrates a pretrained genomic foundation model with hierarchical attention mechanisms to analyze the full protein sequence context of bacterial genomes. GL-HAT is designed to deliver both accurate resistance predictions and interpretable insights into the genomic features associated with resistance using specialized modules. Trained and validated on a comprehensive dataset of bacterial genomes and antimicrobial susceptibility profiles, the model outperformed traditional machine learning method, with an F1-score of 0.845 and AUROC of 0.953. Moreover, GL-HAT could demonstrate the resistance-related contexts in the genome, such as vanA operon and nearby resolvase genes in vancomycin-resistant *Enterococcus* and tet(M) gene adjacent to conjugation proteins in tetracycline-resistant *Streptococcus pneumonia*e. By combining high accuracy with interpretability, GL-HAT offers a practical approach for advancing AMR diagnostics and research, supporting more informed clinical and public health decisions.

## Introduction

Antimicrobial resistance (AMR) presents a persistent challenge to global public health. The spread of resistant pathogens complicates routine clinical care, increases the risk of treatment failure, and contributes to prolonged illness and higher mortality. Beyond individual patient outcomes, AMR accelerates healthcare costs and poses a global public health challenge. Rapid and accurate diagnostic tools are essential to identify resistance profiles early, guide appropriate therapy, and prevent the dissemination of resistant strains^1^.

The clinical standard for AMR detection is culture-based antimicrobial susceptibility testing (AST), including broth dilution and disk diffusion, which directly measure microbial growth in the presence of antibiotics. While considered the gold standard, these methods are labor-intensive and time-consuming^2^. Molecular diagnostic approaches have emerged as alternatives, relying on detection of known resistance genes through nucleic acid amplification or whole-genome sequencing^3^. However, these reference-based methods depend on established catalogs of resistance determinants, missing novel or context-dependent mechanisms.

Comparative genomic analysis provides a route to discover previously unknown resistance factors by contrasting resistant and susceptible isolate groups^4^. This strategy often begins with constructing pangenome presence/absence matrices or summarizing mutation profiles, followed by statistical association to identify candidate genes or variants linked to resistance. Despite its utility, this workflow requires sufficient amount of data of genome-resistance profile pairs for statistical robustness. Also, the definition of meaningful features can be subjective, and resulting models tend to be species-specific and antibiotic-specific, which limits scalability for routine diagnostics.

Computational prediction frameworks have been developed to address these constraints. Classical machine learning methods, such as Random Forest and Support Vector Machines^5^, offer a degree of interpretability but often require separate models for specific species or antibiotic. Moreover, many of these methods still cannot capture the linkages among genes contributing the resistance together^6^. Deep learning models, particularly those based on convolutional neural networks, have demonstrated high predictive performance by capturing non-linear patterns^7,8^. However, limited interpretability often remains a barrier to clinical adoption, and scaling to multi-label predictions across diverse taxa can be computationally demanding.

Recent progress in genomic representation learning enables models to encode long-range genomic context rather than relying solely on isolated gene calls or short motifs^9,10^. Protein language models combined with transformer-based architectures can capture dependencies across operons, mobile genetic elements, and neighborhood structures that often underpin resistance mechanisms. Incorporating such contextual information into predictive frameworks is expected to improve accuracy and provide biologically meaningful interpretability compared to traditional approaches.

In this study, we present a genome-level hierarchical attention transformer (GL-HAT) designed to predict resistance phenotypes across diverse taxa and 53 antibiotics within a single framework. The model leverages a pretrained genomic foundation model (GFM) to capture long-range genomic context and incorporates either of two modules: a max-pooling (MP) module for high-accuracy prediction and a multi-head attention-weighted sum (MH-AWS) module for gene-level interpretability. This approach not only improves predictive performance but also provides interpretable insights into genomic determinants of antimicrobial resistance.

## Materials and Methods

### Dataset collection and preprocessing

Antimicrobial resistance phenotype data were obtained from the PATRIC database^11^, specifically those generated through laboratory methods. To address missing phenotype labels (i.e., Resistant or Susceptible), labels were imputed by identifying entries with identical taxon, antibiotic, experimental method and measurement values that already possessed a label. For cases where the same genome exhibited conflicting phenotype labels for a given antibiotic across multiple experiments, those labels were removed from the dataset, resulting in each genome retaining unique resistance profile labels. Subsequently, the antibiotics with fewer than 500 labeled genomes were excluded from the profile.

Genomes with abnormal sizes, defined as those falling outside the interquartile range (IQR) of [Q1 − 1.5 × IQR, Q3 + 1.5 × IQR] in the corresponding species, were discarded. From the remaining genomes, all unique protein sequences were extracted and clustered using the linclust^12^ with a 50% identity and 90% coverage threshold. Singleton clusters, defined as those with only one representative protein, were excluded.

### Genome clustering and dataset partitioning

For genome clustering, genome similarity was computed using the Jaccard index, defined as the proportion of shared non-singleton protein clusters between two genomes. A similarity threshold of 0.9 was applied to construct a graph where nodes represented genomes and edges connected genomes with the similarity above the cutoff. Clusters were formed by iteratively selecting the genome with the highest number of connections among unassigned genomes as the representative of the cluster and grouping all connected genomes as members. until all genomes with connections were clustered.

For dataset partitioning, cluster representatives were randomly split into a 9:1 ratio and members of each cluster were grouped by resistance profile. To maximize the size of the training set while minimizing bias from overrepresentation of specific clusters, genomes within each profile group in the 90% portion were repeatedly sampled without replacement, with up to 100 genomes per cluster selected. The remaining 10% of cluster representatives were used to construct the validation and test datasets. For each profile group within this portion, one genome with the lowest similarity to the representative was sampled for the validation set. From the remaining genomes in the group, one genome per profile group was sampled for the test set. In the case of singleton clusters, no genome was assigned to the validation set; instead, the single genome was allocated to the test set.

### Feature representation

Each genome was represented as an ordered sequence of proteins, preserving the native genomic arrangement. The amino acid sequence of each protein was embedded using the ESM-2^13^ model (esm2_t33_650M_UR50D), and the mean of the token embeddings was computed to obtain a 1280-dimensional protein-level embedding vector.

### Model architecture

The model was designed as a multi-label classifier to predict resistance profiles across multiple antibiotics for a given genome (Figure 1A). Each genome was represented by its complete set of protein sequences, which were simultaneously input into the model to generate predictions for each antibiotic (Susceptible = 0, Resistant = 1). The protein sequences were embedded using ESM-2^13^ (esm2_t33_650M_UR50D). To capture contextual information across the entire series of proteins within a genome, the model incorporated a hierarchical attention transformer^14^ (HAT) (Figure 1B). HAT consisted of transformer-based segment-wise encoders (SWE) and cross-segment encoders (CSE), as well as segment-wise and cross-segment position embeddings, for capturing local and global context, respectively. The resulting embeddings from the HAT were then passed through classification head, which consisted of two fully connected layers with GeLU^15^ activation followed by layer normalization at between of layers. The GFM was trained while keeping the ESM-2 parameters frozen, allowing the model to learn task-specific representations without fine-tuning the large foundation model.

**Figure 1.**
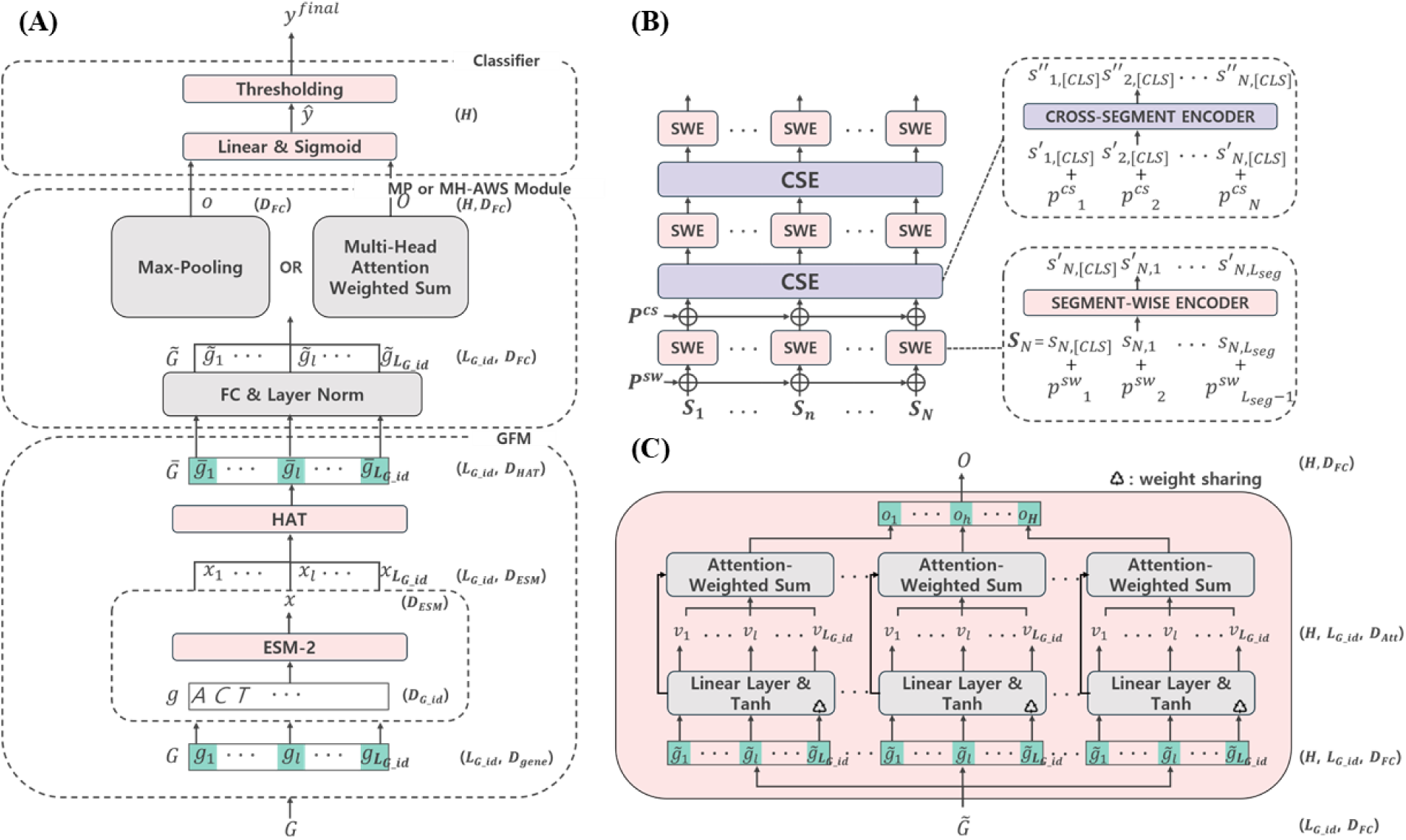
Overview of the GL-HAT model architecture. (A) Overall model workflow, including genomic foundation model (GFM), max pooling (MP) and multi-head attention weighted sum (MH-AWS) module, and classifier. (B) Hierarchical attention transformer (HAT) module, containing layers of segment-wise encoders (SWE) and cross-segment encoders (CSE). (C) MH-AWS module for gene-level attribution and interpretability.

For training GL-HAT, given embedding sequences from pretrained GFM, a fully connected layer followed by layer normalization was applied to project each embedding into an optimal position in the expanded dimensional space. For prediction, two modules were selectively applied: a max-pooling (MP) module, which aggregates gene-wise representations into a single vector for high-accuracy prediction, and a multi-head attention-weighted sum (MH-AWS) module, which produces antibiotic-specific representations for gene-level interpretability (Figure 1C). The final prediction head consisted of a linear layer followed by sigmoid function that produced resistance scores for each antibiotic. Specifically, for the MP module, the linear layer mapped the aggregated single vector to a logit of size equal to the number of antibiotics. For the MH-AWS module, the linear layer mapped each antibiotic-specific representations to a scalar logit value.

### Pretraining genomic context

To enhance predictive performance, the HAT^14^ within the GFM was used to capture contextual information among proteins in each genome. This approach aimed to encode the genomic context as prior knowledge for the resistance prediction. All unique protein sequences found in the dataset were embedded using the ESM-2^13^ model (esm2_t33_650M_UR50D). The resulting embeddings were grouped by k-means clustering using the Faiss^16^ library, with the number of clusters set to 100,000 and default parameters. For each genome, 15% of its proteins were randomly masked, and the model was trained to predict the corresponding k-means cluster of each masked protein. This allowed the model to learn contextual dependencies among proteins based on their co-occurrence patterns. Detailed hyperparameters for MLM-based pretraining are provided in Table S1.

### Model training and evaluation

After the pretraining process, the GFM component was frozen, and the GL-HAT was trained to infer resistance profiles. Detailed hyperparameters for GL-HAT model training are provided in Table S2. After applying a sigmoid activation function to the output scores, a threshold of 0.5 was used to binarize predictions. Evaluation metrics included F1 score, area under the receiver operating characteristic curve (AUROC) and area under the precision-recall curve (AUPRC).

To benchmark the proposed model against conventional approaches, XGBoost^17^ classifier was trained for each antibiotic. The same dataset was used for training, but the input features for XGBoost^17^ were represented as a pangenome presence/absence matrix. This matrix was constructed by clustering all proteins in the dataset using the linclust^12^ with a 90% identity and 80% coverage threshold. Protein clusters found in fewer than 1% of genomes were excluded. The classifiers were implemented using the Python xgboost package with negative log-likelihood evaluation metric and default parameters.

### Component-wise ablation of pretraining and genomic context modules

To assess the contribution of each model component to predictive performance, an ablation analysis was conducted. The evaluation included both the MP and MH-AWS modules, as well as the effects of pretraining the GFM to learn genomic context. The ablation study included GFM with random parameter initialization and a fully connected layer replacing the GFM only for matching the same output dimensions. Each variant was trained and evaluated using the same dataset splits and metrics as the main model. The performance was assessed using F1-score, AUPRC and AUROC.

### Gene-level attribution analysis using MH-AWS

For each antibiotic, the MH-AWS module used multi-head attention to assign gene-level scores, with each head weighing genes differently for that antibiotic. The gene-level scores with respect to a given antibiotic were calculated by taking the exponential of the inner product between the gene representation and the learnable query vector for that antibiotic, then normalizing across all genes in the genome. These scores quantified the relative contribution of each gene in the genome to the predicted resistance for each antibiotic.

To validate the biological relevance of these attributions, we selected genomes from the test dataset that were resistant to vancomycin and tetracycline. For these genomes, we identified known antibiotic resistance genes by running AMRFinderPlus^18^, and for other genes, we utilized annotations provided by the PATRIC database^11^. We then checked the final prediction score for each genome to confirm whether the model’s prediction was correct. Next, we examined the distribution of gene-level attribution scores calculated by the MH-AWS module, and for each genome, we determined which gene corresponded to the peak attribution score for each antibiotic. The product annotation of these peak-scoring genes was then reviewed to assess whether they matched known resistance determinants or other relevant gene products.

## Results

### Experimental settings

The antimicrobial resistance prediction model employed a genomic foundation model based on a hierarchical attention transformer^14^ architecture to capture both local and global genomic context. Each genome was represented as an ordered sequence of protein embeddings generated by ESM-2^13^ and processed through segment-wise and cross-segment encoders. The final prediction was produced using either a max-pooling or multi-head attention weighted sum module, followed by a classifier, as described in Figure 1.

The dataset was constructed by downloading genomes with available antimicrobial susceptibility testing (AST) results from the PATRIC database^11^. Genomes were clustered based on protein composition similarity to ensure unbiased partitioning. As a result, the dataset was split into 59,740 genomes for training, 818 for validation, and 5,061 for testing. Each genome was labeled with available AST results for 53 antibiotics. The number of labels for each antibiotic are provided in Table S3.

Table S4 provides the genus distribution of genomes in each dataset split. *Mycobacterium*, *Streptococcus*, *Escherichia*, *Neisseria*, and *Klebsiella* were among the most abundant genera, with sufficient genomes present in the training, validation and test sets. However, some genera, such as *Morganella*, *Raoultella*, and *Listeria*, had only a small number of genomes, which were found exclusively in the training set.

Table S5 summarizes the number of genomes for each genus-antibiotic pair across the dataset splits. It revealed that certain genera were predominantly associated with specific antibiotics, resulting in differences in data size across pairs. For example, within the genus *Mycobacterium*, data were mainly paired with antibiotics such as rifampin, ethambutol, and isoniazid. Similar patterns were observed for other genera, where a small subset of antibiotics accounted for most of the available data.

### Performance of AMR prediction model

The overall performance of the antimicrobial resistance prediction models was evaluated using F1-score, area under the precision-recall curve (AUPRC), and area under the receiver operating characteristic curve (AUROC) on the full test set (Figure 2A, Table S6). GL-HAT (MP) achieved the highest scores across all metrics, with an F1-score of 0.845, AUPRC of 0.921, and AUROC of 0.953. GL-HAT (MH-AWS) recorded an F1-score of 0.798, AUPRC of 0.883, and AUROC of 0.931, while XGBoost^17^ yielded an F1-score of 0.835, AUPRC of 0.918, and AUROC of 0.950.

**Figure 2.**
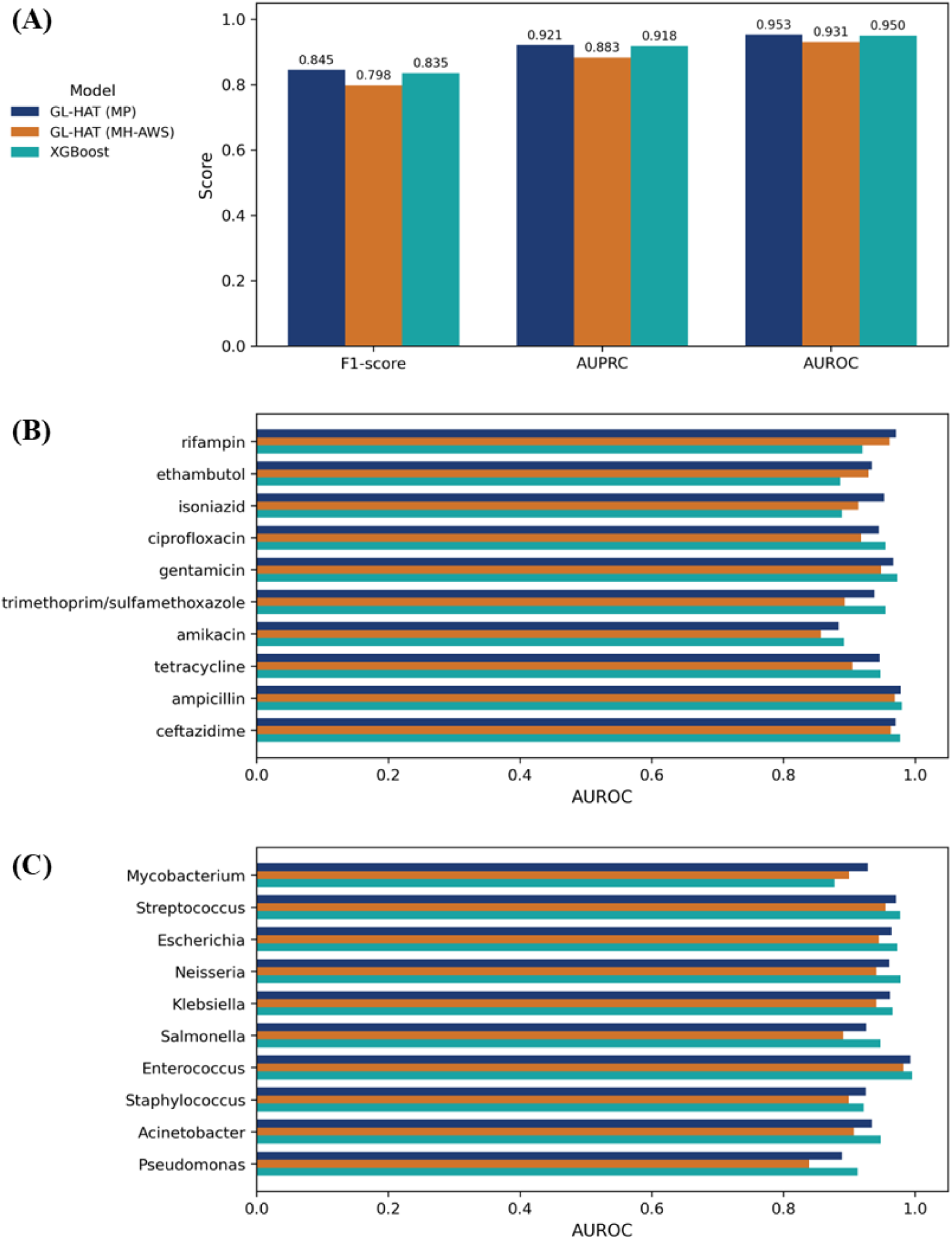
Performance comparison of antimicrobial resistance prediction models. (A) Overall performance metrics (F1-score, AUPRC, AUROC) for GL-HAT (MP), GL-HAT (MH-AWS), and XGBoost on the full test set. (B) AUROC for ten abundant antibiotics. (C) AUROC for ten abundant genera.

Figure 2B shows AUROC values for the ten antibiotics with the largest data sizes. GL-HAT (MP) achieved higher AUROC than the other models for several antibiotics, including rifampin (0.971 for GL-HAT (MP), 0.961 for GL-HAT (MH-AWS), 0.920 for XGBoost) and isoniazid (0.953 for GL-HAT (MP), 0.914 for GL-HAT (MH-AWS), 0.889 for XGBoost). In contrast, XGBoost produced higher AUROC values for some antibiotics. For example, the AUROC for gentamicin was 0.973 with XGBoost, compared to 0.967 with GL-HAT (MP) and 0.948 with GL-HAT (MH-AWS). F1-score, AUPRC and AUROC values for all antibiotics are available in Table S7.

For the ten most prevalent genera in the dataset, the comparison of AUROC values between the models is summarized in Figure 2C. For some genera, GL-HAT (MP) demonstrated higher performance. For example, the AUROC for *Mycobacterium* was 0.928 with GL-HAT (MP), compared to 0.900 with GL-HAT (MH-AWS) and 0.878 with XGBoost. However, XGBoost outperformed the other models for certain genera, such as *Streptococcus*, where XGBoost reached an AUROC of 0.977, whereas GL-HAT (MP) and GL-HAT (MH-AWS) yielded 0.971 and 0.955, respectively. The full set of F1-score, AUPRC, and AUROC values for all genera can be found in Table S8.

Table S9 presents the performance metrics for each genus-antibiotic pair. For pairs with large sample sizes, such as *Mycobacterium* paired with rifampin, ethambutol, and isoniazid, the GL-HAT model usually outperformed XGBoost. This trend was evident in pairs where the training data was abundant, underscoring the advantage of GL-HAT in leveraging rich datasets. In contrast, for pairs with limited data, the performance gap between models was less marked.

### Ablation analysis of model architecture and pretraining effects

To evaluate the contribution of each model component to predictive performance, we conducted an ablation study comparing the MP and MH-AWS modules, as well as the effects of pretraining the GFM to learn genomic context (Table 1).

**Table 1.** Performance comparison of GL-HAT modules and ablation variants. MP: Max-pooling module with pretrained GFM. MP, no pretraining: Max-pooling module without pretrained GFM (random initialization). MP, no GFM: Max-pooling module with a fully connected layer replacing GFM. MH-AWS: Multi-head attention weighted sum module with pretrained GFM. MH-AWS, no pretraining: Multi-head attention weighted sum module without pretrained GFM (random initialization). MH-AWS, no GFM: Multi-head attention weighted sum module with a fully connected layer replacing GFM.

| Model | TP | FN | FP | TN | F1 score | AUPRC | AUROC |
| --- | --- | --- | --- | --- | --- | --- | --- |
| **MP** | 10015 | 2055 | 1615 | 23593 | 0.845 | 0.921 | 0.953 |
| **MP, no pretraining** | 8990 | 3080 | 2514 | 22694 | 0.763 | 0.850 | 0.912 |
| **MP, no GFM** | 9403 | 2667 | 1666 | 23542 | 0.813 | 0.900 | 0.940 |
| **MH-AWS** | 9488 | 2582 | 2210 | 22998 | 0.798 | 0.883 | 0.931 |
| **MH-AWS, no pretraining** | 9680 | 2390 | 2111 | 23097 | 0.811 | 0.896 | 0.939 |
| **MH-AWS, no GFM** | 9417 | 2653 | 2166 | 23042 | 0.796 | 0.882 | 0.932 |

The MP module consistently demonstrated higher predictive performance than the MH-AWS module. The MP module with pretrained GFM achieved an F1-score of 0.845, AUPRC of 0.921, and AUROC of 0.953. Removing pretraining led to a marked reduction in all metrics, with the F1-score dropping to 0.763, AUPRC to 0.850, and AUROC to 0.912. When the GFM was excluded and replaced by a fully connected layer, the MP module yielded intermediate results, with an F1-score of 0.813, AUPRC of 0.900, and AUROC of 0.940.

The MH-AWS module exhibited a distinct pattern. The highest performance for MH-AWS was observed when pretraining was omitted, resulting in an F1-score of 0.811, AUPRC of 0.896, and AUROC of 0.939. In contrast, the configuration with pretrained GFM produced slightly lower values, with an F1-score of 0.798, AUPRC of 0.883, and AUROC of 0.931. Excluding the GFM led to similar decreases, with an F1-score of 0.796, AUPRC of 0.882, and AUROC of 0.932.

### Identification of resistance-related genomic contexts

Despite the lower performance, the interpretation head with the MH-AWS module could represent genomic contexts related to the antibiotic resistance conserved in the pathogen genomes. Rather than randomly distributed across the genome, attribution scores were concentrated within specific genomic contexts that were associated with the resistance label. Functional annotation of the top-scoring loci revealed that the genes with peak scores often corresponded to known resistance genes or to those related to mobile genetic elements.

To investigate the genomic contexts underlying vancomycin resistance, we analyzed the test set genomes labeled as resistant and annotated with AMRFinderPlus^18^. As shown in Figure 3A, the majority of vancomycin-resistant genomes harbored the vanA operon (n=55), a smaller subset carried vanB (n=8), and two genomes contained both operons, with no other vancomycin operons detected.

**Figure 3.**
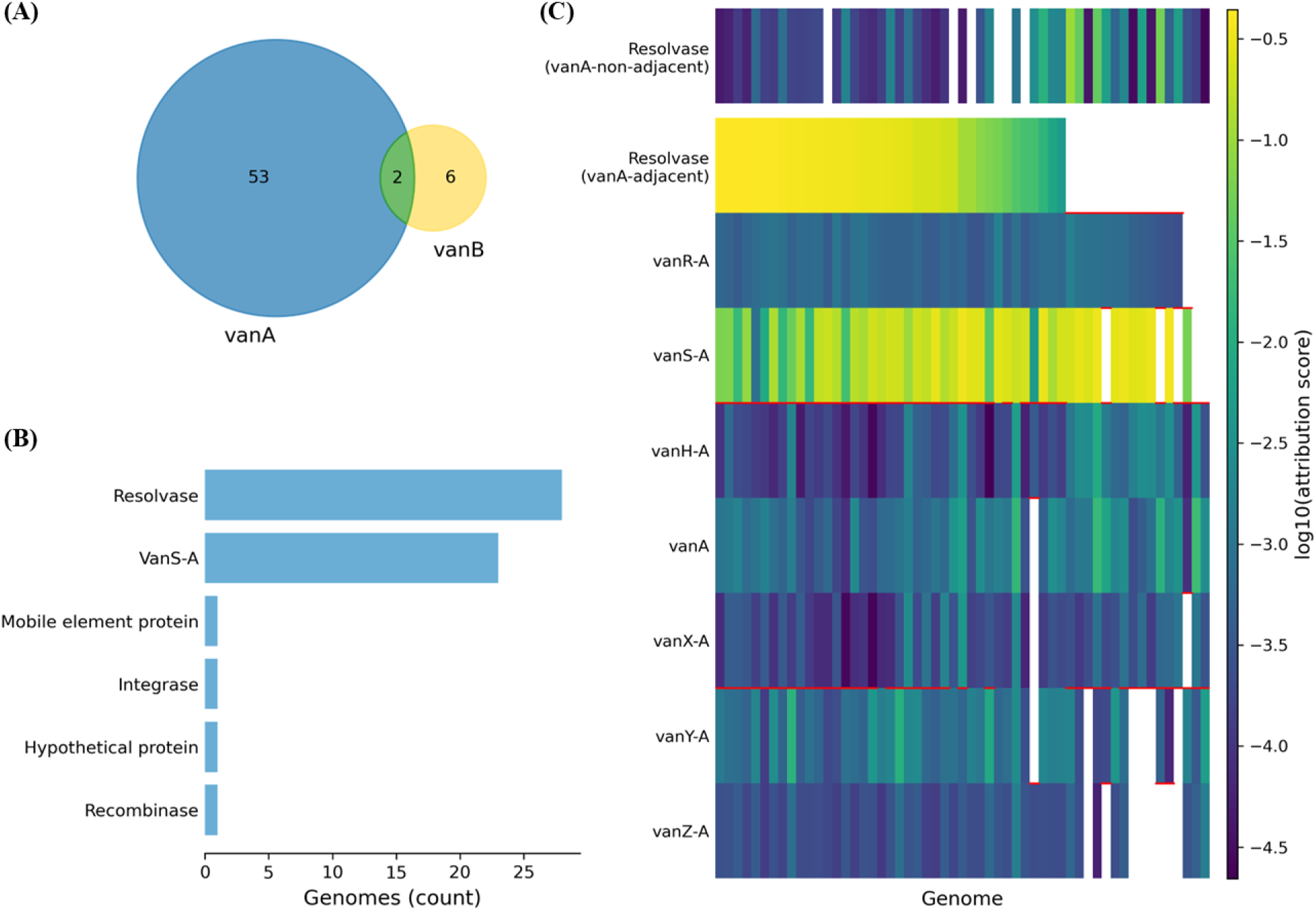
Gene-level analysis by attribution scores for vancomycin-resistant genomes. (A) Venn diagram showing counts of genomes including vanA and vanB operons (55 vanA, 8 vanB, 2 both; total 61). (B) Peak-attribution products among vanA-positive genomes (n=55): resolvase and vanS dominated the highest-attribution positions, with minor contributions from mobile element protein, integrase, hypothetical protein and recombinase. (C) Heatmap of gene-level attribution across the vanA operon, vanA-adjacent resolvase and non-adjacent resolvase, with log-scaled scores; missing genes appear in white. Horizontal red separators indicate boundaries where neighboring operon genes were located on different contigs within a genome.

In the case of the distribution of attribution scores across vanA-positive genomes, the scores showed a steep rank-order decay (Table S10). The median ratio of the second-highest to the highest score was approximately 0.32, while the third-highest fell below 0.05, indicating that the top attribution typically stood out against background weights. We then examined which gene products were most frequently assigned the highest attribution scores by the MH-AWS module among vanA-positive genomes (Figure 3B). More than half of the genomes (28/55, 50.9%) exhibited peak attribution at a resolvase gene, while vanS was the most highly attributed gene in 23 genomes (41.8%). Other products, including mobile element protein, integrase, hypothetical protein and recombinase, were rarely assigned the highest score.

To clarify the relationship between the resolvase genes with the highest scores and vancomycin resistance, we visualized the gene-level attribution scores for the vanA operon and the adjacent resolvase across all vanA-positive vancomycin resistant genomes in the test set (Figure 3C). The heatmap showed that, when a resolvase gene was present immediately upstream of vanA operon, this gene consistently received the high attribution score among the operon and its neighborhood. In contrast, resolvase genes located elsewhere in the genome, even when multiple copies were present and their scores aggregated, generally exhibited low attribution values. Occasional high scores were observed only in genomes lacking a vanA-adjacent resolvase. Additionally, vanS showed high attribution scores across many genomes, regardless of whether the resolvase gene or other operon genes were present on the same contig. Even in cases where the vanA operon was fragmented across multiple contigs, vanS maintained high attribution scores.

In contrast to the vancomycin analysis, which in our test set comprised *Enterococcus* genomes predominantly carrying vanA, tetracycline resistance spanned multiple species and displayed heterogeneous resistance gene calls and product annotations at the highest-attribution positions. We stratified tetracycline-resistant genomes by species and summarized AMRFinderPlus detections together with the product of the gene receiving the top MH-AWS attribution in each genome. Figure 4A represents species-specific associations, including tet(M) in *Streptococcus pneumoniae* with a conjugation-related protein, tet(B) in the *Acinetobacter calcoaceticus*/*baumannii* complex with hypothetical proteins.

**Figure 4.**
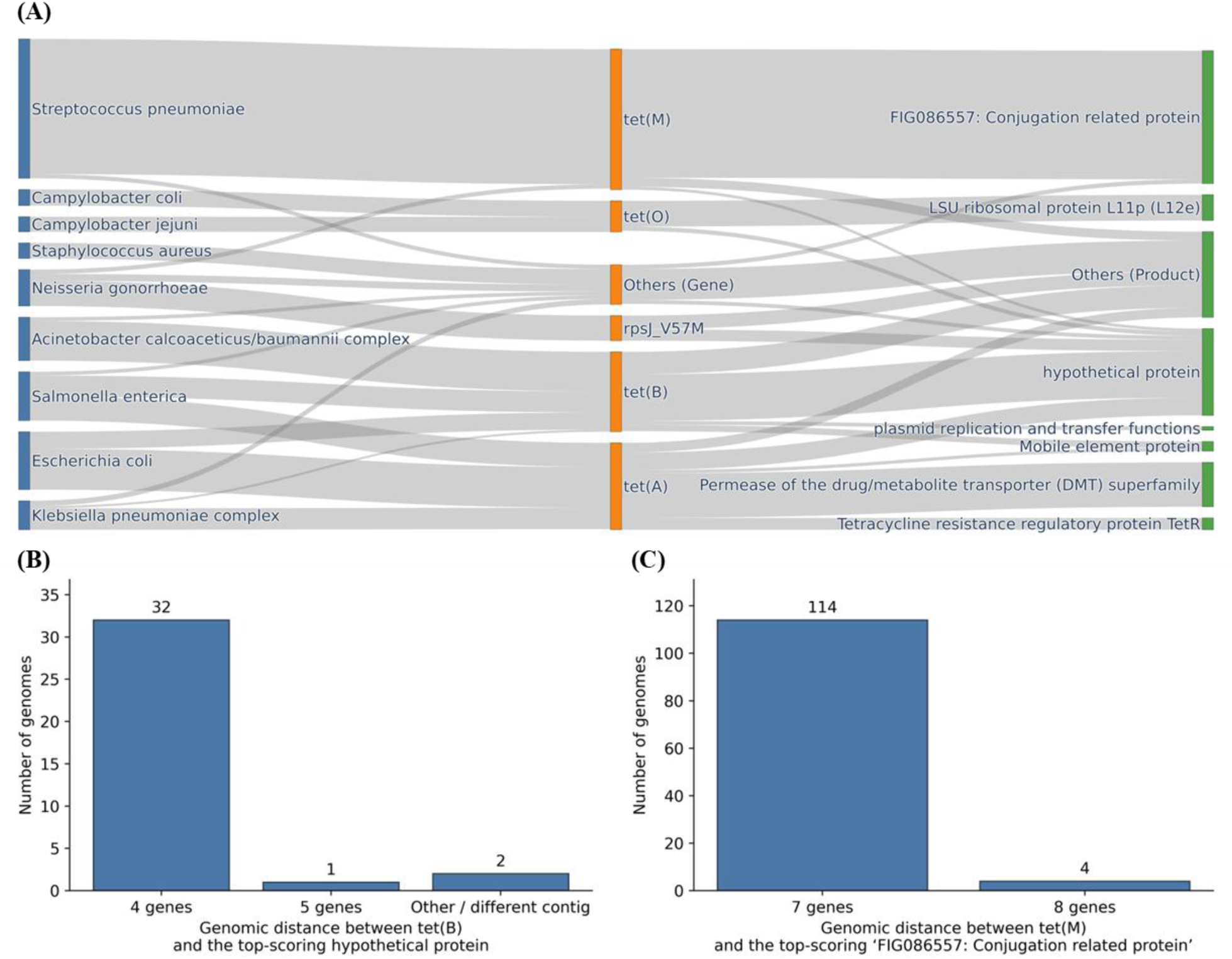
Species-specific tetracycline resistance contexts captured by MH-AWS attribution. (A) Sankey diagram linking species (left), AMRFinderPlus-detected tetracycline resistance genes (middle), and the product annotation of the gene with the highest attribution per genome (right). (B) Distribution of gene-order distance between tet(B) and the top-scoring hypothetical protein within *Acinetobacter calcoaceticus*/*baumannii* complex genomes that both harbor tet(B) and exhibit a hypothetical protein as the highest-attribution position (n = 35). “Other / different contig” category comprises distances except 4 or 5, and cross-contig pairs. (C) Distribution of gene-order distance between tet(M) and the top-scoring Conjugation related protein in *Streptococcus pneumoniae* (n = 118).

Within the tetracycline-resistant *Acinetobacter calcoaceticus*/*baumannii* complex, 53/55 genomes in the test set were correctly predicted. Of these, 37 carried tet(B) by AMRFinderPlus, and 37 had a hypothetical protein as the highest-attribution product; 35 genomes satisfied both criteria. For these 35 genomes, we measured the gene-order distance between tet(B) and the top-scoring hypothetical protein on assembled contigs. A prominent mode at four genes was observed (32/35), with one genome at five genes and two genomes either at other separations or on different contigs (Figure 4B).

In *Streptococcus pneumoniae*, 130/145 tetracycline-resistant genomes were correctly predicted; 124 carried tet(M) and 124 had FIG086557: Conjugation related protein as the highest-attribution product, with 118 genomes meeting both conditions. The gene-order distance between tet(M) and the top-scoring conjugation-related protein was highly conserved: seven genes in 114/118 genomes and eight genes in 4/118 genomes (Figure 4C).

## Discussion

This study introduces a genome-level hierarchical attention transformer (GL-HAT) for antimicrobial resistance (AMR) prediction across diverse bacterial taxa and 53 antibiotics. By integrating a pretrained genomic foundation module (GFM) with hierarchical attention mechanisms, the model achieved superior predictive performance compared to conventional machine learning approach. The max-pooling (MP) module delivered the highest accuracy, while the multi-head attention-weighted sum (MH-AWS) module provided gene-level interpretability, enabling insights into genomic determinants of resistance.

Machine learning models such as Random Forest and XGBoost^17^ have been widely applied for AMR prediction but typically require separate classifiers for each antibiotic, limiting scalability and failing to capture correlations among resistance phenotypes. Ensemble-based multi-label methods partially address this issue but introduce complexity and computational overhead. Deep learning approaches using convolutional neural networks have demonstrated improved accuracy but often lack interpretability and generalizability across species. GL-HAT overcomes these limitations by modeling long-range genomic context and multi-label dependencies within a single framework, reducing the need for multiple models and improving performance on large-scale datasets.

Beyond predictive accuracy, interpretability is critical for clinical adoption and biological discovery. Attention-based attribution in GL-HAT consistently highlighted genomic regions associated with known resistance mechanisms, such as vancomycin operons. For vancomycin, GL-HAT attribution maps frequently peaked at resolvase and vanS in vanA-positive genomes, rather than only the classical vanA gene. This is consistent with the biology of the Tn1546 transposon, where resolvase is conserved as the context related to the transposition of vanA operon^19–21^.

For tetracycline resistance, the attribution score distribution in *Streptococcus pneumoniae* consistently highlighted the genomic context of the tet(M) gene and conjugation-related protein with a fixed distance from it, which are typically embedded within the conjugative transposons of Tn916 family^22^. These findings suggest that the model captures biologically meaningful patterns through attribution scores rather than simply relying on the existence of resistance gene itself. Moreover, this attribution score analysis can be expanded to the identification and annotation of previously unknown context related to the resistance, like hypothetical proteins close to tet(B) in *Acinetobacter calcoaceticus*/*baumannii* complex genomes.

Ablation studies confirmed the importance of pretraining and contextual modeling for MP module. Incorporating a pretrained GFM improved performance compared to models without pretraining or genomic context, underscoring the value of transfer learning in bioinformatics. By leveraging masked language modeling on protein sequences, the model learned dependencies that extend beyond individual genes, capturing operon-level and neighborhood-level patterns relevant to resistance. Meanwhile, for the MH-AWS module, the use of a pretrained GFM showed the reduction in accuracy compared to the model without pretraining. This potentially reflects the fact that the attention mechanism in MH-AWS inherently learns task-specific context during antibiotic-level representation aggregation, reducing the incremental benefit of external pretraining.

Despite these strengths of GL-HAT, several limitations require consideration. Performance varied across taxa and antibiotics, with lower accuracy observed for rare genera and antibiotics with highly imbalanced labels. The model may be sensitive to genome assembly quality and contig fragmentation, which could disrupt contextual representation. Additionally, attention-based interpretability, while informative, does not establish causality and should be complemented by experimental validation.

It is also important to note that, outside of *Mycobacterium* and other combinations with abundant training data, XGBoost^17^ often outperformed GL-HAT for many genus-antibiotic pairs. This is partly attributable to XGBoost approach of training separate classifiers for each antibiotic, which can be advantageous when data are limited or highly imbalanced. However, this also highlights the critical need for continued data supplementation and balanced sampling to fully utilize the benefits of context-aware deep learning models. As more comprehensive and diverse datasets become available, the strengths of GL-HAT in capturing genomic context and biological relationships are expected to become increasingly valuable for robust resistance prediction.

Future work will focus on expanding the model to additional taxa and antibiotics, as well as adapting it for metagenomic samples to support environmental and clinical surveillance. Improving inference speed and reducing memory footprint will be essential for integration into real-time clinical workflows.

Combining attention-based interpretation with other attribution methods and incorporating experimental validation will strengthen the biological relevance of model outputs.

## Code Availability

The codes for the GL-HAT model described in this study is available at the following Github repository: https://github.com/cjbs-bi/GL-HAT_paper.

